# SARS-CoV-2 transmission chains from genetic data: a Danish case study

**DOI:** 10.1101/2020.05.29.123612

**Authors:** Andreas Bluhm, Matthias Christandl, Fulvio Gesmundo, Frederik Ravn Klausen, Laura Mančinska, Vincent Steffan, Daniel Stilck França, Albert H. Werner

**Author notes:** These authors contributed equally to this work.

## Abstract

**Background:** The COVID-19 pandemic caused by the SARS-CoV-2 virus started in China in December 2019 and has since spread globally. Information about the spread of the virus in a country can inform the gradual reopening of a country and help to avoid a second wave of infections. Denmark is currently opening up after a lockdown in mid-March.

**Methods:** We perform a phylogenetic analysis of 742 publicly available Danish SARS-CoV-2 genome sequences and put them into context using sequences from other countries.

**Result:** Our findings are consistent with several introductions of the virus to Denmark from independent sources. We identify several chains of mutations that occurred in Denmark and in at least one case find evidence that the virus spread from Denmark to other countries. A number of the mutations found in Denmark are non-synonymous, and in general there is a considerable variety of strains. The proportions of the most common haplotypes is stable after lockdown.

**Conclusion:** Our work shows how genetic data can be used to identify routes of introduction of a virus into a region and provide alternative means for verifying existing assumptions. For example, our analysis supports the hypothesis that the virus was brought to Denmark by skiers returning from Ischgl. On the other hand, we identify transmission chains suggesting that Denmark was part of a network of countries among which the virus was being transmitted; thus challenging the common narrative that Denmark only got infected from abroad. Our analysis does not indicate that the major haplotypes appearing in Denmark have a different degree of virality. Our methods can be applied to other countries, regions or even highly localised outbreaks. When used in real-time, we believe they can serve to identify transmission events and supplement traditional methods such as contact tracing.

## Introduction

The first cases of the COVID-19 pandemic were reported in the city of Wuhan (China) in December 2019 and a new virus, named SARS-CoV-2, was later identified as its origin [1]. At the time of writing, the pandemic is ongoing and has spread to more than 180 countries [2].

The first case in Europe was reported from France on January 24, 2020 [3]. Italy confirmed the first two cases on January 31 [4]. Austria reported its first cases on February 25 [5]. In March, Europe was the center of the global pandemic with many European countries introducing lockdown measures and travel restrictions. Early on, the ski area of Ischgl in Tyrol, Austria, was identified as a transmission hot-spot by some countries, so Iceland already declared it a risk area on March 5 [6]. Quarantine measures in Ischgl, however, were only imposed on March 13 [7].

The first case in Denmark was confirmed on February 27, after having returned from skiing holidays in Northern Italy on February 24 [8]. The second case was confirmed on February 28 after having entered Denmark from Italy on February 15 [9]. The number of cases kept increasing, and, on March 8, when two cases at Rysensteen Gymnasium were confirmed [10], there was increased suspicion of community transmission within Denmark.

During this first phase of the pandemic in Denmark, there existed travel warnings to high risk areas. On March 2, Denmark advised against all travel to Northern Italy [11]. On March 10, Denmark additionally advised against travel to the Austrian state of Tyrol, as the country had seen many infected skiers returning from the ski village of Ischgl [12].

On March 11, the Danish prime minister Mette Frederiksen announced a lockdown, which happened in several stages and included closures of borders and schools [13]. Overall, the measures were not as severe as in some other European countries. On April 6, the prime minister announced that a first phase of reopening would start from April 14 [14]. The country has opened up further since.

As of May 26, there were 11,428 confirmed infections and 563 deaths in Denmark in connection with the disease [15]. From March 12, only people with serious symptoms and people in risk groups were tested. Since April 1, the number of tests has been increased [16].

In this work, we study all the publicly available genome sequences of the SARS-CoV-2 virus from Denmark as of May 26, and compare them to sequences from abroad. See S2 File for a list of sequences and their labs of origin. We use the mutations in the genomic data to identify transmission chains. We focus on chains highlighting the introduction of the virus to Denmark, its transmission within Denmark, and its spread to other countries.

## Materials and methods

We use publicly available sequenced genome of the SARS-CoV-2 virus in order to identify mutations for the purpose of finding transmission chains. In the following, we describe how we obtained and analyzed the sequences. For a flow chart of this process, see Fig 1.

**Fig 1.**
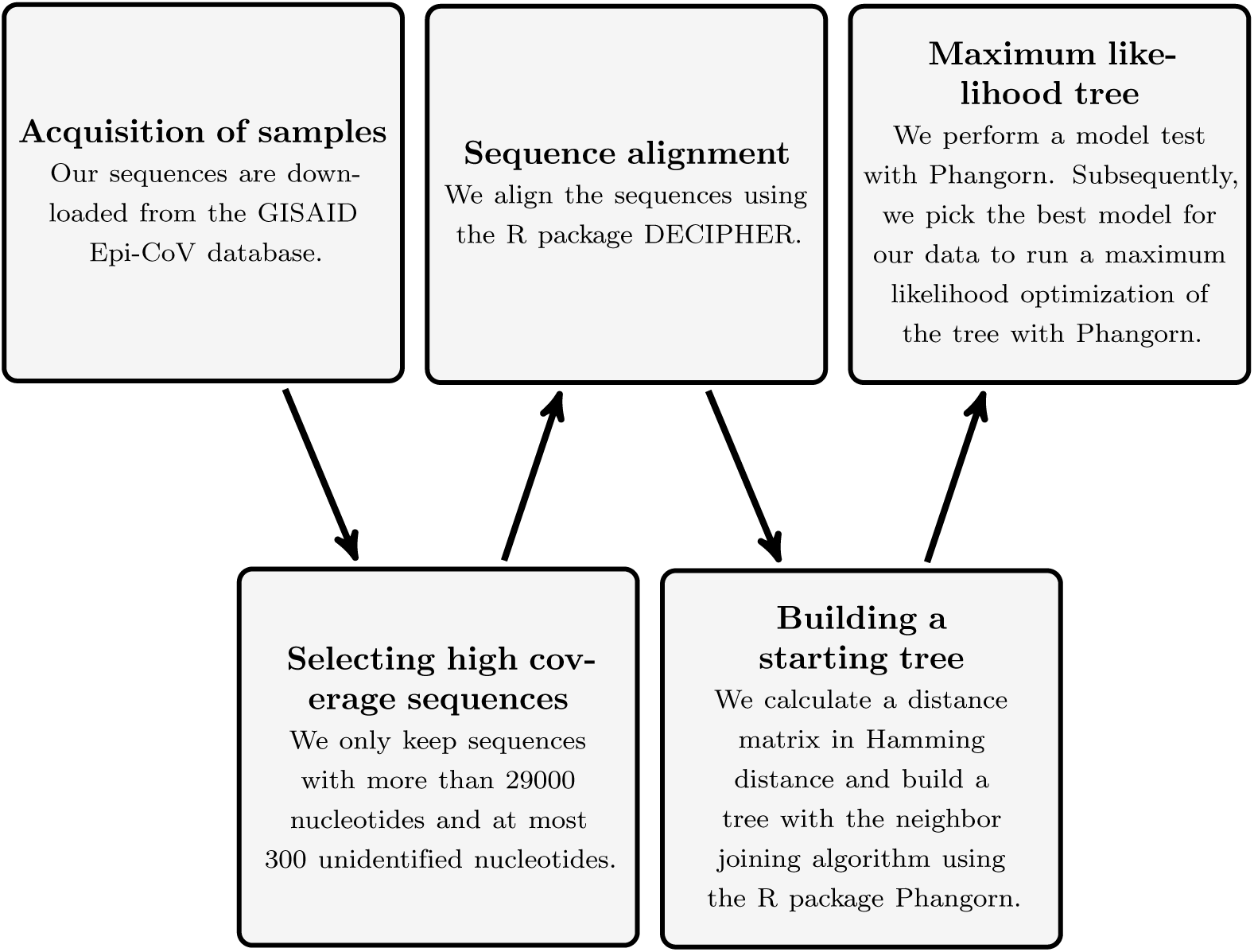
The tree building process as a flow chart.

### Acquisition of samples

Sequences were downloaded from the GISAID EpiCoV database [17, 18] on May 26, 2020, including 742 Danish sequences. See the supplementary file S2 File for a full list of sequences including their origins. From the available sequences, we selected those which we deemed of high quality and used them for our analysis. Specifically, we only consider sequences with at least 29,000 nucleotides having at most 300 unidentified nucleotides (N’s). Moreover, we only consider sequences that originate from a human host. After these steps, we were left with 582 Danish sequences, which we focus on in our analysis.

### Sequence alignment

We perform multiple sequence alignment (MSA) using the R-package *DECIPHER* [19, 20]. We apply the function AlignSeqs from this package to our data until convergence is achieved.

### Tree inference

The tree topology was inferred using the R-package *phangorn* [21]. From the aligned data, we build a distance matrix in Hamming distance using the function dist.hamming. In the next step, we build a phylogenetic tree from this distance matrix using the neighbor joining algorithm (function NJ in *phangorn*).

This tree is then used as the starting tree for a maximum likelihood optimization. We run the modelTest function to identify the most suitable probabilistic model for the given tree and data. Based on the outcome of this step, we then generate a maximum likelihood tree using the optim.pml function. To further validate the generated tree, we perform bootstrapping using the bootstrap.pml function with 2000 samples for each tree.

Tree visualization and annotation were done with the R-package *ggtree* [22]. We plot the trees ignoring branch lengths to focus on the tree topology.

Most of our analysis uses R [23] and except for the R-packages mentioned in the previous sections, we also use *stringr, dplyr, ggplot2* and *ape* [24]. The code we used can be found at https://github.com/qmath/phylo_qmath.

### Haplotypes and rooting conventions

We root our trees with respect to the reference sequence NC-045512.2 (SARS-CoV-2 isolate Wuhan-Hu-1), which is also the reference for Nextstrain [25] and for [26]. We choose naming conventions for haplotypes in accordance with [26] based on a naming convention from Nextstrain [25], which seems most suitable for our purposes. In Table 1, we list the mutations corresponding to the names we will use, following [26, Table S3]. In addition, we list their names in the more recent Nextstrain convention [27] and the clades from the pangolin system that they are included in [28]. Finally, we list the corresponding amino acid changes and in which genes they can be found. See [29, Fig 1] for an overview of the SARS-CoV-2 genome.

**Table 1.** Naming of different haplotypes.

| [26] | mutations | new Nextstrain | pang. | amino acid change |
| --- | --- | --- | --- | --- |
| A2 | C241T, C3037T, A23403G | 19A/C241T/C3037T/A23403G | B | D614G in S |
| A2a | A2+C14408T | 19A/C241T/C3037T.20A | B.1 | A2+P4715L in Orf1ab |
| A2a1 | A2a+GGG28881AAC | 19A/C241T/C3037T.20A.20B | B.1.1 | A2a+R203K+G204R in N |
| A2a2 | A2a+G25563T | 19A/C241T/C3037T.20A/G25563T | B.1 | A2a+Q57H in Orf3a |
| A2a2a | A2a2+C1059T | 19A/C241T/C3037T.20A.20C | B.1 | A2a2+T265I in Orf1a |

List of relevant haplotypes with their definition in terms of mutations as well as their new Nextstrain label. We note that both the reference string used and all the haplotypes listed have the four mutations specified as haplotype A in [26]. We therefore do not list those. The second to last column is the label for the currently identified pangolin lineage clades they are included in. The last column gives the corresponding amino acid changes and the genes they occur in.

To get an overview of haplotypes and mutations prevalent in Denmark, we identify positions where sufficiently many of the analyzed sequences exhibit a substitution or a deletion as compared to the other sequences in the data set. For a better overview and readability we choose different thresholds depending on the context. We analyze the co-occurrence of the new mutations with previously identified haplotypes from [26] and with each other in the entire worldwide data set.

## Results

In this section, we review our results. After a general overview of the mutations over time, we study three different types of mutations in more detail: First, we consider mutations which were present in some region of the world and appeared in Denmark at some point. Second, we look at chains of mutations which only appear in Denmark. Finally, we look at mutations for which a Danish origin is predominant. In the Discussion section, we analyze the spread of the virus to, within and from Denmark in light of these mutations. The mutations identified by this haplotype analysis were cross-validated via bootstrapping for the corresponding phylogenetic trees, with resulting bootstrap values consistent with the number of mutations defining the different clades. Let us illustrate this using the tree built for the sequences containing the mutation C15842A, shown in the section Chain of mutations starting at C15842A. In this tree we observe a bootstrap value larger than 86 for clades defined by two additional mutations. On the other hand, for clades defined by a single mutation, we observe bootstrap values around 65, as expected.

### Distribution of mutations over time in Denmark

The most common haplotypes in Denmark are A2a2a and A2a1. Fig 2 shows in the top panel the relative weight of those haplotypes over time with a seven-day rolling average. In the lower panel we plot the seven-day rolling average of the number of sequences. We observe a larger fraction of A2a1, which is associated to Italy, before the lockdown. From the onset of the lockdown, the fractions stay rather stable in time with A2a2a, which we discuss in detail below, making up around 70%. A lower number of sequences may be interpreted as a larger error bar on the haplotype percentages, making the haplotype distribution in time consistent with constant proportions.

**Fig 2.**
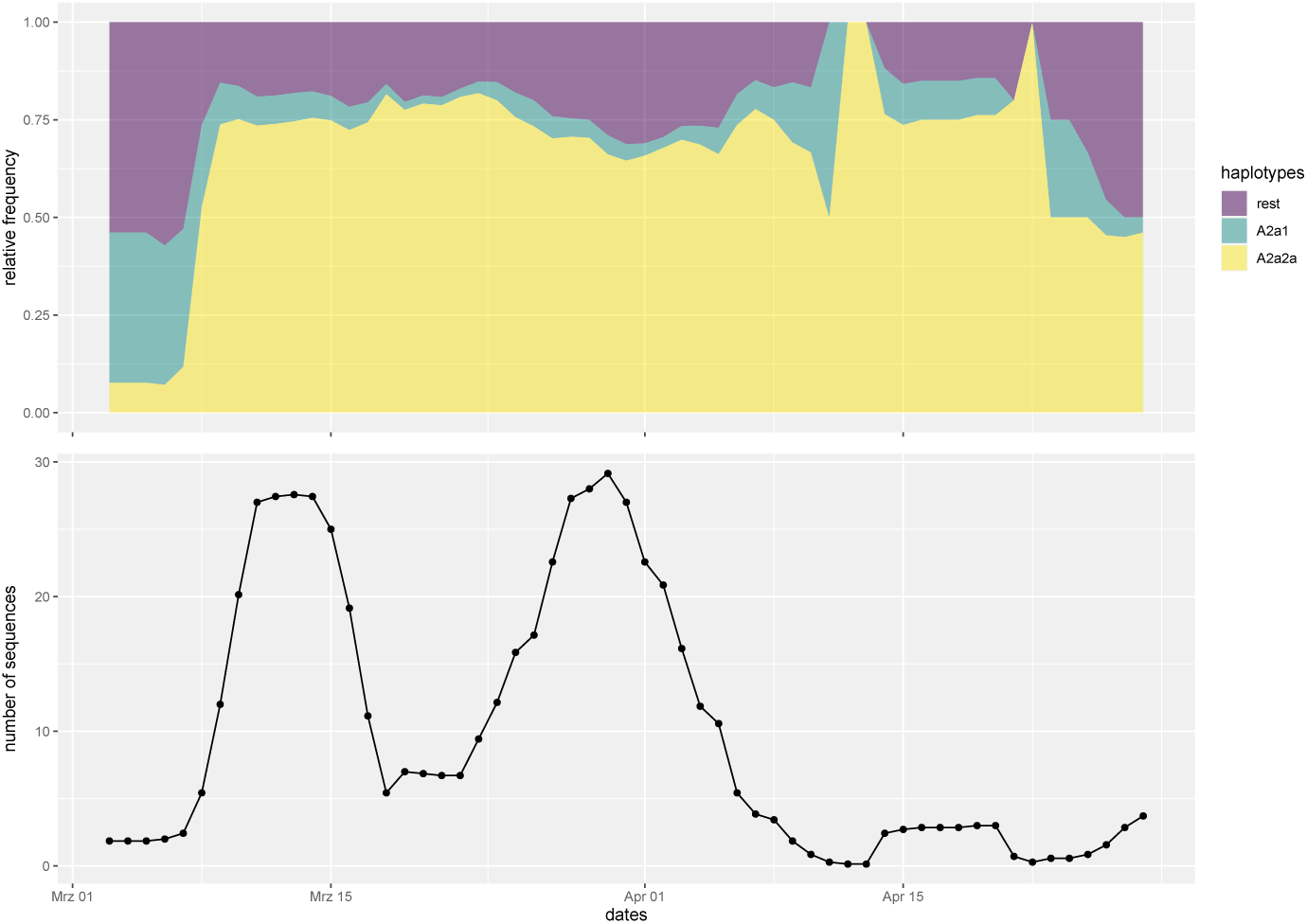
Haplotypes over time. Top panel: relative frequencies of major haplotypes in Denmark over time; bottom panel: total number of sequences over time. Both graphs are based on seven-day rolling averages.

### Mutations from other regions appearing in Denmark

In the following, we first discuss the most common Danish haplotypes in relation to their appearance in other countries. We subsequently discuss specific examples of less prevalent mutations in Denmark that are present to a larger extent in other countries. For an overview we refer to Fig S1 in S1 File.

#### A2a2a: A common mutation in Denmark and Ischgl

Approximately 70% of the available Danish sequences have haplotype A2a2a. This makes it the most common haplotype in our Danish data with 405 out of 582 sequences. In our complete data set, we see that 4343 out of 20239 sequences have this haplotype (see Table S1 in S1 File), with sequences originating from the US, Denmark, the UK, Australia, France and other countries. This haplotype was already reported in [26] where the authors point out that travelers from Austria had the haplotype A2a2 together with the mutation C1059T which is the definition of A2a2a. The haplotype A2a2 corresponds to an amino acid change Q57H in Orf3a as compared to A2a and the haplotype A2a2a corresponds to an amino acid change T265I in Orf1a as compared to A2a2 (see also Table 1). Both mutations have already been studied in [30].

In the following we first give evidence that some of the sequences with A2a2a originate from the skiing area of Ischgl in the region of Tyrol, Austria, by identifying specific transmission chains. We will then argue that most, but likely not all of this haplotype comes from that area.

In order to identify a specific transmission chain, we will look at mutations that occur in addition to A2a2a. Consider therefore mutation A6825C which corresponds to the amino acid change N2187T in Orf1a and which is seen in nine sequences that have A2a2a worldwide. These include six Danish ones, while the others are from Austria, Norway and Scotland. The Norwegian sequence can be traced with metadata to Austria. The Norwegian sequence is dated to March 9, which makes it likely that this mutation has been present in Austria prior to that date. It is therefore consistent that the Danish sequences, the first of which also is dated to March 9, originate from Austria. Since the Austrian sequence is from Ischgl, a spread from Ischgl, a tourist skiing destination, seems likely.

Similarly, the mutation G15380T (corresponding to S5039L in Orf1a) appears with haplotype A2a2a in 32 sequences worldwide. Among those 32 are 16 of Danish origin. Of the remaining ones with A2a2a, there are eight Austrian sequences. All of them stem from the region of Tyrol and in particular six are from Ischgl.

In order to argue that most Danish sequences with A2a2a originate from Austria, we first observe that the ratio of sequences with A2a2 versus A2a2a is close to 1 in Germany, Denmark, Norway, Austria, Iceland, Sweden and Switzerland what regards European countries, and lower in other European countries such as the UK, France and the Netherlands from which significant travel to Denmark would be expected. Since the number of Danish sequences with A2a2a is high even when restricting to the time around the onset of the lockdown, there must have been multiple introductions of A2a2a to Denmark, and it therefore seems unlikely that this could have happened from a country with a much different ratio than that of Denmark.

Tourism from European countries to Alpine ski resorts around February and March would provide a natural travel route for the virus, in particular to Denmark, Sweden, Norway and Germany. For Iceland, this assumption is supported by the travel information collected in [26].

The sequences from Switzerland have no travel histories, but location data shows that the Swiss sequences with A2a2a are mainly spread across the German-speaking part and that they are in particular not concentrated at one location. This makes it unlikely that there was a hotspot in a Swiss ski resort.

In contrast, the Austrian sequences can be mainly attributed to the skiing region of Ischgl. We refer to the Austrian data presented in Fig S2 of S1 File, where one sees that the haplotype A2a2a is mostly present in sequences from the ski village of Ischgl in the region of Tyrol, Austria, and the adjacent region of Vorarlberg.

Accordingly, it seems very plausible that Ischgl was indeed a hotspot for the transmission of the haplotype A2a2a to the aforementioned countries.

The Norwegian sequences have travel metadata and give further supporting evidence for this infection route. Here, three out of twelve sequences with the haplotype A2a2a also have recent travel history to Austria (the others having unknown travel history).This is shown in Fig S3 in S1 File. The Icelandic study [26] also associated the haplotype A2a2a with travel to Austria.

However, due to the abundance of the haplotype A2a2a in the world, it is likely that some portion of the sequences with the haplotype A2a2a are not part of transmission chains through Ischgl. As an example we discuss A2a2a + G24368T in Subsection S1.4 of S1 File, which we identify as likely originating from the UK.

#### A2a1: A common mutation in Denmark and Italy

We see that the haplotype A2a1 appears 38 times in Denmark. Moreover, 36 sequences with haplotype A2a1 were found in the early targeted testing group (January 31-March 15) in the Icelandic study [26, Table 2]. Out of these, 29 had a travel history from Italy and three from Austria. Furthermore, the earliest Danish sequence (dated February 26) is from when there was only one confirmed case in Denmark. As reported in the news, this case has travel history to an Italian ski-area. It also has haplotype A2a1.

#### The triple deletion ATGA1605A with coincident mutation T514C

Both in the Danish and the worldwide data sets we observe sequences with a triple deletion at sites 1606–1608 (ATGA1605A) which is sometimes coincident with a substitution T514C (identified as A6 in [26, Table S3]). The triple deletion corresponds to the triple deletion ATGA1604A identified as haplotype A9 in [26, Table S3]. Note that [26, Table S3] places it at position 1604 rather than 1605, which seems to be a typo. The CoV-GLUE database confirms the deletion at the nucleotides where we find it [31].

Most of the sequences in our data set with the triple deletion ATGA1605A but without the substitution T514C are from the UK (293 out of 346). In contrast, most of the sequences with both the deletion ATGA1605A and the substitution T514C are from the Netherlands (98 out of 138). In addition, the earliest of the sequences with both ATGA1605A and T514C are from the Netherlands as well. Therefore, we conclude it to be likely that the triple deletion originated in the UK and then spread to the Netherlands, where it picked up the mutation T514C. In Denmark, we observe nine sequences with ATGA1605A, two of which additionally have T514C. These latter two Danish sequences are likely of Dutch origin. See Fig 3 for an illustration. The mutation T514C is not shown in Fig S1 of S1 File, since it appears only twice in the Danish data. However, the sequences in question are those at the very bottom, with numbers EPI_ISL_444828|2020–03–11 and EPI_ISL_429295|2020–03–13.

**Fig 3.**
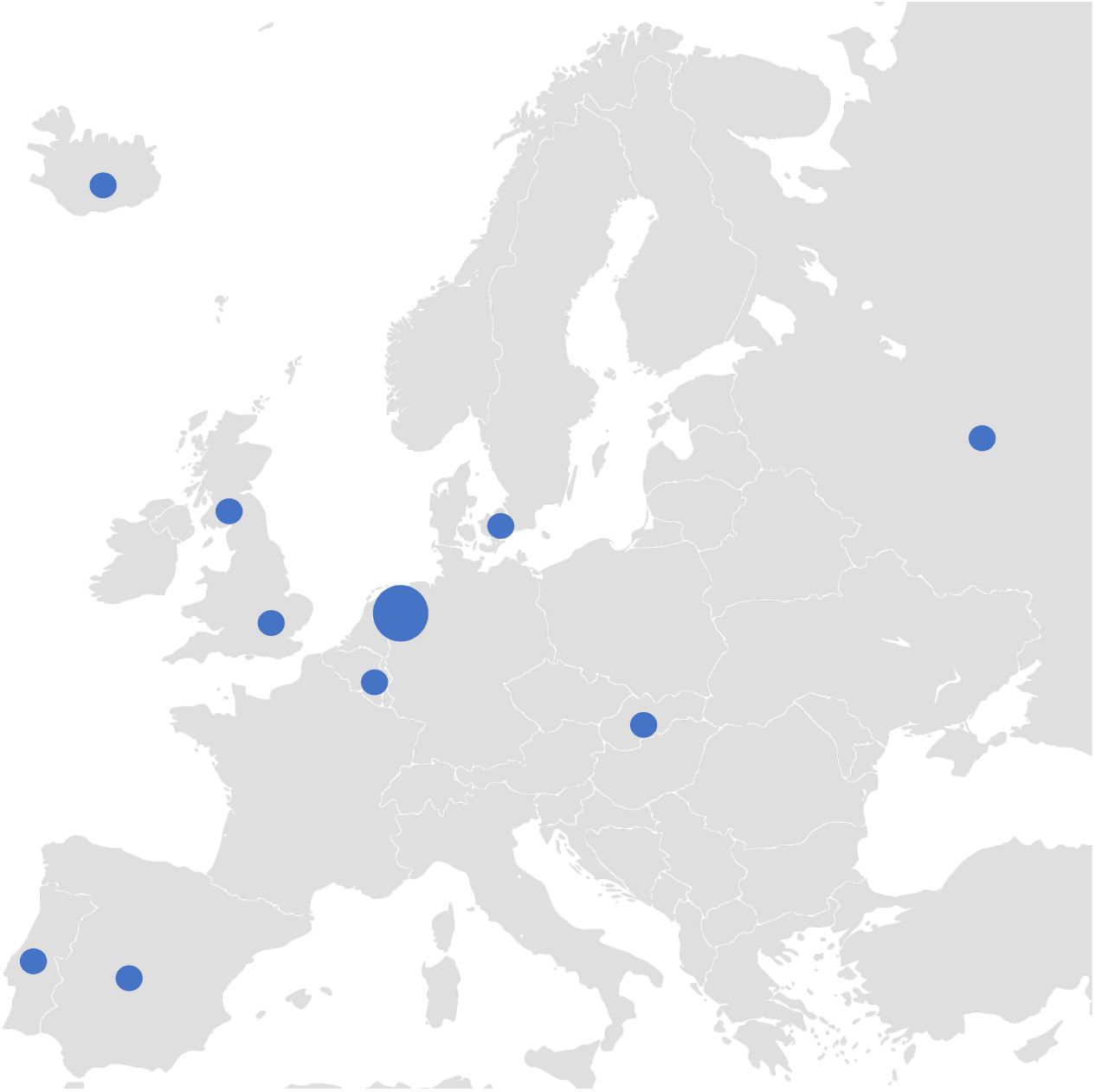
Appearance of the strain with haplotype ATGA1605A and T514C. According to a maximum likelihood analysis, it likely originated from the Netherlands and spread to other European countries, including Denmark.

#### Chains of mutations starting in Denmark

Now, we turn to chains of mutations which occurred inside Denmark. From Fig S1 in S1 File, one identifies several chains of mutations. Here we report two of the most pronounced.

#### Chain of mutations starting at C15842A

The phylogenetic tree with an overview of the associated haplotypes for this mutation can be found in Fig 4. There are 20 sequences with the mutation C15842A and the haplotype A2a2a worldwide and they are all of Danish origin.

**Fig 4.**
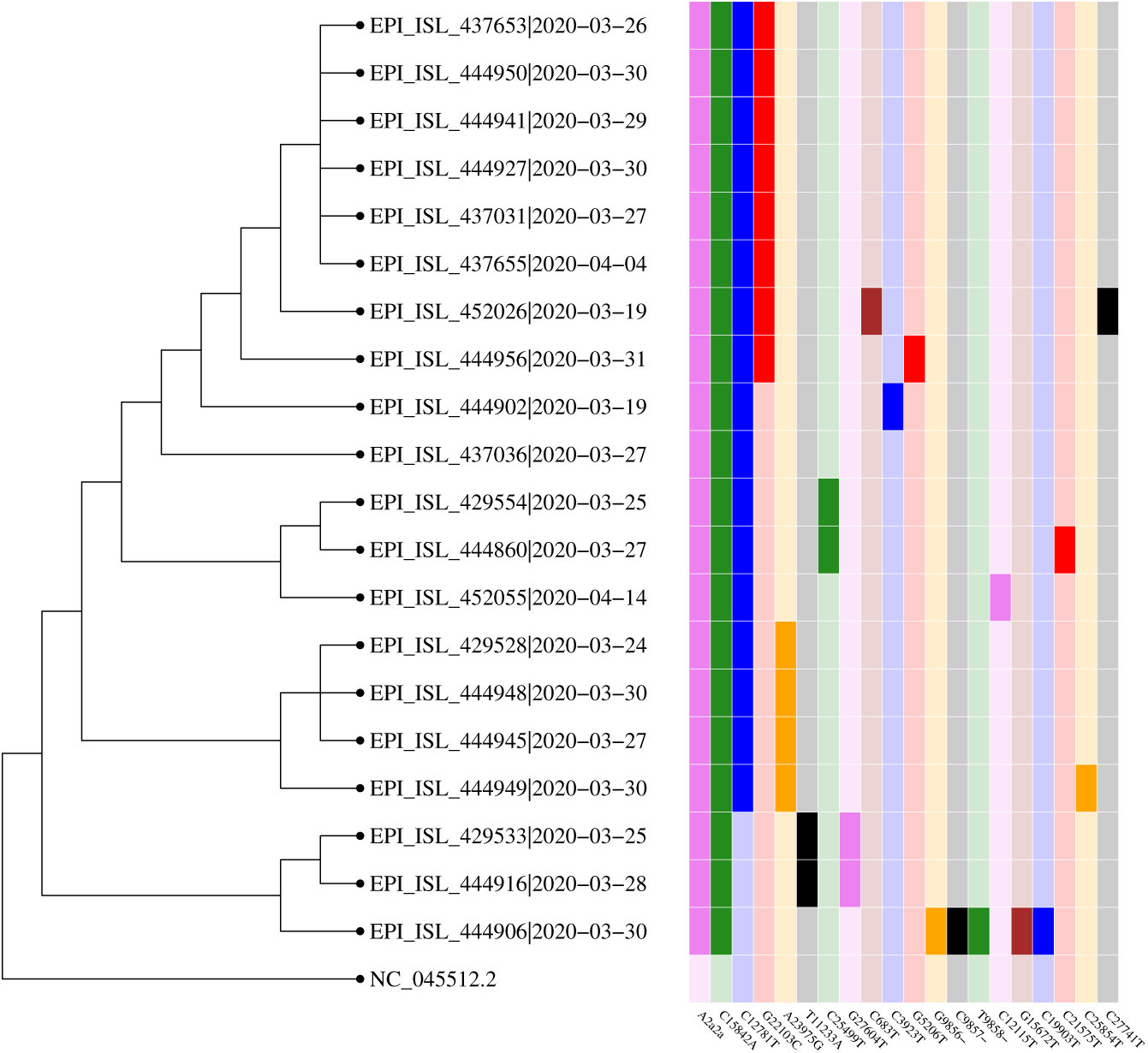
Phylogenetic tree for sequences containing mutation C15842A. The second mutation shown is C15842A, followed by C12781T. After that, it trifurcates into G22103C, A23975G and C25499T.

From the 20 (all Danish) sequences with A2a2a and C15842A, there are 17 which also have the mutation C12781T. Furthermore, of the sequences that have both the mutations C15842A (T5193N in Orf1a) and C12781T (synonymous), there are eight which in addition have the non-synonymous mutation G22103C (G181R in the spike protein). Another four sequences have the mutation A23975G instead and finally, there are two which have C25499T. Some of the previously mentioned sequences have additional mutations. The longest chain of mutations appearing at least twice has length three (not counting the mutations composing the haplotype A2a2a; see Fig 5).

**Fig 5.**
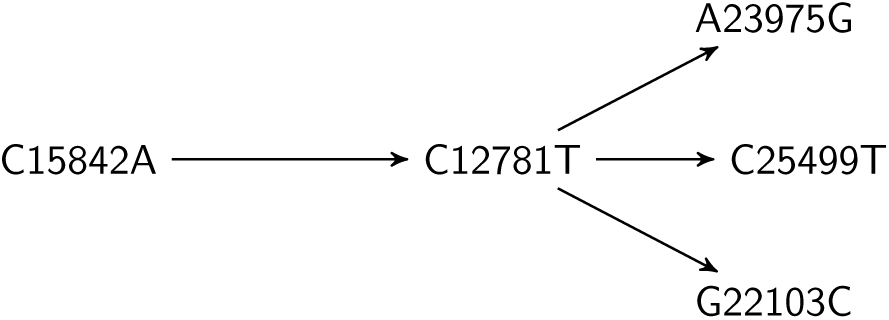
Chain of mutations starting at C15842A.

#### Chain of mutations starting at C1302T

The Danish sequences with haplotype A2a2a show frequently the mutation C1302T. It is non-synonymous and corresponds to amino acid change T346I in Orf1a. We will argue that this mutation originated in Denmark and that it mutated further in Denmark as well as spread to other countries.

In order to see that this mutation spread further from Denmark, note that worldwide there are 115 sequences with the mutation C1302T co-occurrent with the haplotype A2a2a, 103 of which are Danish. The remaining ones are Latvian (1), Icelandic (5) and Swedish (6). The travel histories of the five Icelandic sequences show that two have traveled to Denmark (as first reported in [26]), while the other cases do not contain travel information. Of the six Swedish sequences, one is from Uppsala dated to March 12 while the five others are from Norrbotten (in the north of Sweden) dated from March 24 until April 2. The earliest Danish sequence with C1302T is from March 3. Whereas our analysis does not completely exclude that the virus spread from Sweden or Latvia to Denmark, we believe that the earlier date of the Danish sequence, together with the high abundance in Denmark, makes it very likely that it originated in Denmark and further spread from Denmark.

See also Fig 6 for an illustration of the international presence of the mutation. In order to see that the strain A2a2a+C1302T further mutated in Denmark we inspect the corresponding clade of the Danish tree in Fig S1 of S1 File. Ten of the sequences have C11074T (a combination which is not found outside of Denmark). Of these, six have the mutation C29095T. Three of them moreover have the mutation A9280G, whereas two have the mutation C619T (this is not visible in the plot due to a threshold of three when displaying mutations). Of the ones with mutation A9280G, two have a mutation at C7164T. Some of the sequences have additional single mutations. We have thus identified the Danish chains of mutations in Fig 7.

**Fig 6.**
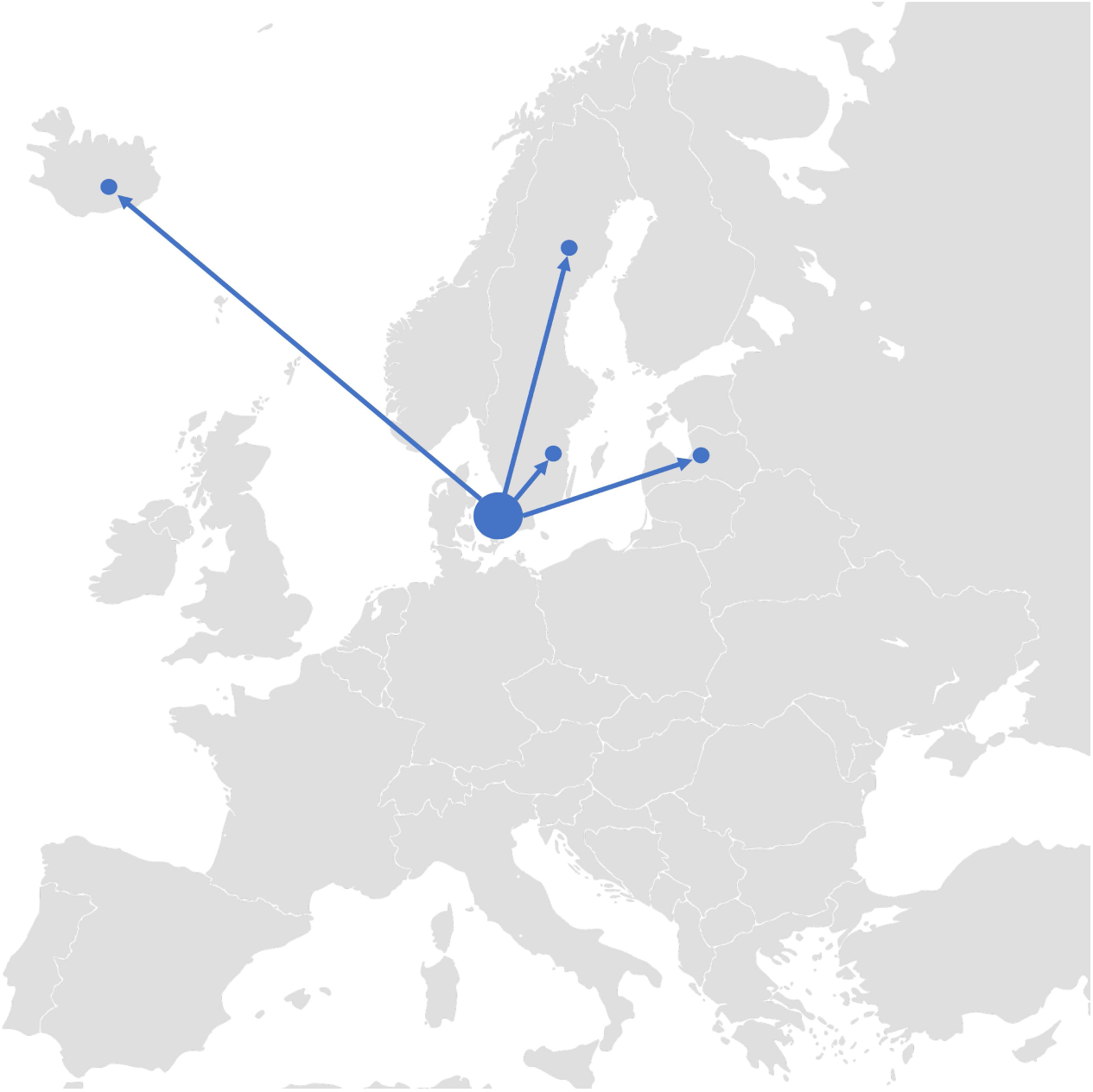
Spread of the strain with haplotype C1302T. The figure shows the likely spread of the strain with haplotype C1302T from Denmark to other Northern European countries. Within Denmark, it also mutated further.

**Fig 7.**
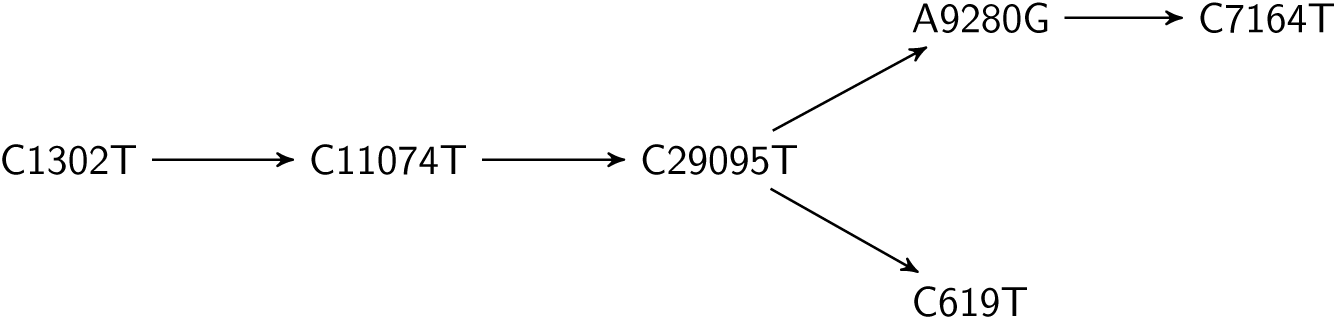
Chains of mutations starting C1302T.

## Discussion

### Haplotypes over time

During the initial period of the introduction of the virus to Denmark from different sources the percentages of the different major haplotypes change: From a larger proportion of haplotype A2a1 associated to Italy to a 70% proportion A2a2a associated to Ischgl. After lockdown, however, we do not observe a significant change in the proportions anymore. Therefore, we find no evidence for different virality. We find no clear pattern in individual mutations of the strains appearing in Denmark either. Therefore, we suspect that they are consistent with random mutation events.

### Introduction to Denmark

Our phylogenetic analysis shows that around 70% of the Danish sequences have haplotype A2a2a. By comparing the distribution of haplotypes across different countries, by utilizing date information, and, for some international sequences, also location data as well as travel histories, we conclude that the majority of the Danish sequences with A2a2a originate directly or indirectly from Ischgl. This is illustrated with examples of specific transmission chains from Ischgl. Our observation that a large proportion of Danish sequences originate Ischgl is not unexpected given the public knowledge of travel histories [32]. Our analysis, however, can be regarded as an independent cross-check of this existing narrative.

The remaining portion of the sequences is consistent with multiple entries from other countries, among them Italy, the UK and the Netherlands. We have illustrated this with example transmission chains which can be associated to those countries. In the case of Italy, this is based on the haplotype A2a1, in the case of the UK, it is a specific mutation on top of haplotype A2a2a and in the case of the Netherlands, it is a mutation in addition to a well-known triple deletion. These conclusions are consistent with the testing results in mid-March [33].

From this point of view, our genomic analysis cross-validates the public statements and supports findings in the Iceland study [26] that indicate that Ischgl was a hotspot earlier than widely recognized.

### Transmission chains inside Denmark

We have listed all mutations that appear at least three times inside Denmark in Table S2 in S1 File and also plotted them in Fig S1 of S1 File. We note that many more Danish transmission chains can be identified from Fig S1 of S1 File.

In the results, we discussed two particularly pronounced chains of mutations based on this plot and Fig 4. For the two chains described in the results we conclude that they are transmission chains that happened inside Denmark. We have chosen these two since these mutation chains only co-occur with the haplotype A2a2a in Denmark (except of the first mutation C1302T). This shows clearly how one can track the virus mutating as it spreads inside Denmark. The longest chain that we conclude happened inside Denmark is five mutations long and consists of the mutations

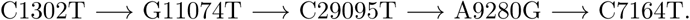

These mutations took place in a period from before March 15 to before April 14 based on the dating of the sequences. The average mutation rate was estimated in [34] as 10^−4^ nucleotides/genome/year and it would be interesting to cross check the mutation rate with the number of mutations in the Danish data.

### Transmission out of Denmark

For the mutation C1302T, based on its high prevalence in Denmark compared to the rest of the world together with the travel histories of the Icelandic cases, we conclude that it has appeared in Denmark and spread from there to Sweden, Latvia and to Iceland. Some reservations remain since the Swedish data in GISAID is very limited with only 163 sequences as of May 26. Further, the transmission chain described shows how the virus has spread extensively within Denmark and mutated at least four times after that.

Hence we see indications that the virus has mutated several times inside Denmark and spread from Denmark, as illustrated in Fig 6. We have listed and discussed the most common mutations. Some of the mutations we see seem to have occurred independently elsewhere, in particular in the UK, which has a large number of sequences in GISAID.

### Outlook

The conclusions above are based on a rather large number of high quality Danish sequences with date information as well as on sequences from other countries which in part have more metadata. Even though we do not have firm knowledge of the representativity of the Danish sequences, we can assert that they cover the entire time from the first identified case up to May 9. In order to confirm the analysis of the proportion of haplotypes seen, it would be important to supplement this with information about the representativity of the analysed data set or obtain access to a more representative sample. At the same time, our analysis also shows how to effectively incorporate metadata (date, country of origin or travel history) in such an analysis.

The presented ability to identify specific transmission chains and routes of introduction from SARS-CoV-2 genomic data displays its potential for understanding the spread of the virus in a population. Although here we present a case study for Denmark, a similar analysis could be carried out for outbreaks in other countries, regions or even smaller units such as hospitals. If the genomic data is available in real-time, such an analysis can inform mitigation measures even during an ongoing outbreak, for instance supplementing traditional methods such as contact tracing.

## Supporting information

Acknowledgement tables for GISAID

Extra tables and figures

## Supporting information

**S1 File. Discussion of additional Danish mutations and supplementary tables and phylogenetic trees**. (PDF)

**S2 File. GISAID acknowledgements**. List of sequences from GISAID used and their submitting laboratories. (PDF)

## Acknowledgments

We thank Judith Gottwein, Anders Krogh, Jakob Sture Madsen and Carsten Wiuf for valuable comments. We acknowledge financial support from VILLUM FONDEN via the QMATH Centre of Excellence (Grant no. 10059). AHW thanks the VILLUM FONDEN for its support with a Villum Young Investigator Grant (Grant No. 25452). We would like to thank everyone submitting sequenced genome data to GISAID, in particular Statens Serum Institut, with whom we shared an earlier version of this manuscript, and Mads Albertsen’s lab. A full list of the contributors can be found in S2 File.

## Note

During completion of this work, we became aware of concurrent work, which was announced here [35].

## Notes

### Competing Interest Statement

The authors have declared no competing interest.

