## Supplementary material for "SARS-CoV-2 transmission chains from genetic data: a Danish case study": Extra tables and figures

### S1 Discussion on additional common Danish mutations

In this section, we discuss some of the most common Danish mutations (with respect to the Wuhan reference) in addition to the ones already discussed in the main text. We will use another haplotype from [26], namely A8. This haplotype corresponds to G1440A and G2891A with respect to the Wuhan reference sequence. We refer to Tables S1 and S2 for a summary of mutations occurring at least three times in the Danish data.

**S1.1 Mutation C7011T.** Worldwide there are 64 sequences with the mutation C7011T, of which 38 exhibit haplotype A8. Among these 38 are 13 Danish sequences. We suspect that the mutation occurred independently at least twice. One occurrence seems to be from the UK; the other cluster (the one co-occurrent with A8) is less clear. The cluster with the Danish sequences contains five Austrian sequences. The Austrian sequences are from Innsbruck, Tyrol, Ischgl, Vienna and St. Anton. This means that four out of five are from the region of Tyrol, but they are not concentrated in Ischgl. In the overview of the Austrian haplotypes in Fig S2 the haplotype A8 is missing in three of the Austrian sequences, but they all have at least one of the two mutations constituting A8. Some of the sequences have an ambiguity character here, so it could be that most Austrian sequences have A8.

**S1.2 Mutations T8788C and G26951A.** Worldwide there are 55 sequences with the mutation T8788C and 56 with mutation G26951A. Almost all are Danish. In Fig S1 one can also see that those two mutations are co-occurring and that they are in the beginning of several transmission chains.

**S1.3 Mutation C7834T.** Worldwide there are 73 sequences with the mutation C7834T, of which 66 are with haplotype A2a. Among those are the 60 Danish ones so the mutation C7834T constitutes one most abundant Danish mutations and in Fig S1 one can see that they are starting transmission chains.

**S1.4 Mutation G24368T.** Worldwide there are 163 sequences with the mutation G24368T, of which 156 are with haplotype A2a2a. Among those are the five Danish ones (all with haplotype A2a2a). The majority of the others are from the UK (89), Sweden (50), the Netherlands (4), Iceland (3). The three Icelandic samples all exhibit a travel history connected to the UK. The mutation is non-synonymous and is located in the spike protein. The mutation G24368T seems closely tied to the UK and we note that compared to the total number of sequences, a much larger fraction of the Swedish sequences is of this type (50/163) compared to the corresponding fraction of Danish sequences (5/582). This indicates that Sweden might have been infected through the UK to a larger extend than Denmark. Another point of interest is that G24368T is co-occurring with haplotype A2a2a. This is a specific example that the link between haplotype A2a2a and Austria is non-exclusive.

### S2 Supplementary tables

| Total | World 20239 | Denmark 582 | Sweden 163 | Norway 33 | France 375 | Italy 84 | UK 6668 | Netherlands 489 | Austria 247 | Germany 173 |
|---|---|---|---|---|---|---|---|---|---|---|
| A | 20220 | 580 | 162 | 32 | 374 | 83 | 6645 | 488 | 246 | 172 |
| A1a | 1418 | 0 | 1 | 6 | 11 | 7 | 901 | 50 | 4 | 6 |
| A1a1 | 1372 | 0 | 1 | 1 | 9 | 0 | 898 | 50 | 4 | 6 |
| A1a1a | 546 | 0 | 0 | 0 | 2 | 0 | 271 | 45 | 3 | 1 |
| A1a1a1 | 42 | 0 | 0 | 0 | 0 | 0 | 0 | 0 | 0 | 0 |
| A1a1a2 | 14 | 0 | 0 | 0 | 0 | 0 | 0 | 0 | 0 | 0 |
| A1a1a3 | 8 | 0 | 0 | 0 | 0 | 0 | 0 | 0 | 0 | 0 |
| A1a1a4 | 84 | 0 | 0 | 0 | 0 | 0 | 59 | 0 | 0 | 0 |
| A1a1a5 | 27 | 0 | 0 | 0 | 1 | 0 | 15 | 0 | 0 | 0 |
| A1a1b | 645 | 0 | 1 | 0 | 7 | 0 | 523 | 2 | 0 | 5 |
| A1a2 | 0 | 0 | 0 | 0 | 0 | 0 | 0 | 0 | 0 | 0 |
| A2 | 13900 | 553 | 153 | 25 | 332 | 76 | 5024 | 314 | 211 | 141 |
| A2a | 13867 | 552 | 153 | 25 | 332 | 76 | 5024 | 314 | 211 | 126 |
| A2a1 | 4363 | 38 | 43 | 3 | 26 | 36 | 2690 | 110 | 96 | 50 |
| A2a1a | 363 | 13 | 1 | 0 | 2 | 1 | 85 | 83 | 34 | 17 |
| A2a1a1 | 27 | 0 | 0 | 0 | 0 | 0 | 0 | 0 | 0 | 0 |
| A2a1a2 | 28 | 0 | 0 | 0 | 0 | 0 | 7 | 4 | 0 | 8 |
| A2a1a3 | 12 | 1 | 0 | 0 | 0 | 0 | 8 | 0 | 0 | 0 |
| A2a1b | 349 | 4 | 1 | 0 | 0 | 0 | 280 | 2 | 2 | 2 |
| A2a1c | 230 | 0 | 1 | 0 | 1 | 0 | 153 | 5 | 4 | 0 |
| A2a1d | 6 | 0 | 0 | 0 | 0 | 0 | 0 | 0 | 0 | 0 |
| A2a2 | 5343 | 410 | 70 | 14 | 185 | 0 | 421 | 52 | 66 | 44 |
| A2a2a | 4343 | 405 | 66 | 12 | 112 | 0 | 279 | 34 | 63 | 44 |
| A2a3 | 943 | 13 | 3 | 1 | 1 | 2 | 425 | 19 | 22 | 2 |
| A2a3a | 82 | 0 | 0 | 0 | 0 | 0 | 0 | 0 | 0 | 0 |
| A2a4 | 13 | 0 | 0 | 0 | 0 | 0 | 0 | 0 | 0 | 0 |
| A2a5 | 195 | 3 | 3 | 6 | 1 | 5 | 63 | 5 | 0 | 0 |
| A2a6 | 104 | 4 | 0 | 0 | 0 | 0 | 22 | 48 | 5 | 2 |
| A2a7 | 432 | 0 | 0 | 0 | 0 | 0 | 409 | 0 | 0 | 0 |
| A2a8 | 513 | 1 | 0 | 0 | 110 | 0 | 43 | 6 | 1 | 2 |
| A2a9 | 136 | 1 | 1 | 0 | 2 | 6 | 51 | 7 | 2 | 6 |
| A2a10 | 125 | 0 | 0 | 0 | 0 | 0 | 92 | 0 | 0 | 0 |
| A2a11 | 23 | 0 | 0 | 0 | 0 | 0 | 2 | 19 | 0 | 0 |
| A2a12 | 26 | 0 | 0 | 0 | 0 | 1 | 7 | 5 | 0 | 0 |
| A2a13 | 149 | 2 | 0 | 0 | 0 | 0 | 131 | 0 | 0 | 0 |
| A3 | 162 | 1 | 1 | 1 | 0 | 0 | 2 | 2 | 0 | 1 |
| A6 | 158 | 2 | 0 | 0 | 0 | 0 | 19 | 99 | 3 | 1 |
| A7 | 17 | 0 | 0 | 0 | 0 | 0 | 4 | 0 | 0 | 0 |
| A8 | 310 | 16 | 3 | 0 | 0 | 0 | 189 | 6 | 5 | 17 |
| A9 | 73 | 0 | 0 | 0 | 0 | 0 | 63 | 0 | 0 | 0 |
| A10 | 3 | 0 | 0 | 0 | 0 | 0 | 0 | 0 | 0 | 0 |
| B | 2173 | 1 | 2 | 0 | 2 | 0 | 72 | 7 | 5 | 2 |
| B1 | 1299 | 0 | 0 | 0 | 0 | 0 | 5 | 0 | 0 | 0 |
| B1a | 1275 | 0 | 0 | 0 | 0 | 0 | 5 | 0 | 0 | 0 |
| B1a1 | 1249 | 0 | 0 | 0 | 0 | 0 | 5 | 0 | 0 | 0 |
| B1a1a | 124 | 0 | 0 | 0 | 0 | 0 | 0 | 0 | 0 | 0 |
| B1a1a1 | 13 | 0 | 0 | 0 | 0 | 0 | 0 | 0 | 0 | 0 |
| B1a1a1a | 10 | 0 | 0 | 0 | 0 | 0 | 0 | 0 | 0 | 0 |
| B1a1a1b | 1 | 0 | 0 | 0 | 0 | 0 | 0 | 0 | 0 | 0 |
| B2 | 46 | 0 | 0 | 0 | 0 | 0 | 0 | 0 | 0 | 1 |
| B4 | 98 | 0 | 2 | 0 | 0 | 0 | 0 | 3 | 0 | 0 |

TABLE S1. **Summary of haplotypes for selected European countries as per May 26, 2020.** We include the haplotypes described in [26].

| Total | World 20239 | Denmark 582 | Sweden 163 | Norway 33 | France 375 | Italy 84 | UK 6668 | Netherlands 489 | Austria 247 | Germany 173 |
|---|---|---|---|---|---|---|---|---|---|---|
| C1302T | 115 | 103 | 6 | 0 | 0 | 0 | 0 | 0 | 0 | 0 |
| C7834T | 73 | 60 | 0 | 0 | 0 | 0 | 4 | 0 | 0 | 0 |
| G26951A | 56 | 54 | 0 | 0 | 0 | 0 | 2 | 0 | 0 | 0 |
| T8788C | 55 | 53 | 0 | 0 | 0 | 0 | 2 | 0 | 0 | 0 |
| C15842A | 20 | 20 | 0 | 0 | 0 | 0 | 0 | 0 | 0 | 0 |
| C12781T | 37 | 17 | 0 | 0 | 0 | 0 | 0 | 1 | 0 | 0 |
| G15380T | 40 | 16 | 0 | 1 | 1 | 0 | 5 | 0 | 8 | 2 |
| C7011T | 64 | 13 | 2 | 0 | 0 | 0 | 28 | 1 | 5 | 0 |
| C21707T | 51 | 11 | 0 | 0 | 0 | 0 | 4 | 0 | 0 | 0 |
| C11074T | 48 | 11 | 0 | 0 | 0 | 0 | 19 | 1 | 0 | 0 |
| C29200T | 41 | 8 | 0 | 0 | 0 | 0 | 21 | 0 | 0 | 0 |
| G22103C | 8 | 8 | 0 | 0 | 0 | 0 | 0 | 0 | 0 | 0 |
| C28253A | 8 | 7 | 0 | 0 | 0 | 0 | 0 | 0 | 0 | 0 |
| A6825C | 10 | 6 | 0 | 1 | 0 | 0 | 1 | 1 | 1 | 0 |
| C3743T | 8 | 6 | 0 | 0 | 0 | 0 | 1 | 0 | 0 | 0 |
| C16887T | 72 | 5 | 1 | 0 | 3 | 0 | 32 | 0 | 0 | 0 |
| G19086T | 13 | 6 | 0 | 0 | 0 | 0 | 3 | 0 | 0 | 1 |
| G24368T | 163 | 5 | 50 | 0 | 1 | 0 | 95 | 4 | 0 | 0 |
| T99C | 4 | 4 | 0 | 0 | 0 | 0 | 0 | 0 | 0 | 0 |
| C218T | 8 | 4 | 0 | 0 | 0 | 0 | 1 | 0 | 0 | 0 |
| A518G | 7 | 3 | 0 | 0 | 0 | 0 | 0 | 0 | 0 | 0 |
| T658C | 26 | 4 | 0 | 0 | 0 | 0 | 1 | 4 | 0 | 2 |
| C4002T | 174 | 4 | 1 | 0 | 0 | 0 | 118 | 2 | 2 | 0 |
| A4548G | 3 | 3 | 0 | 0 | 0 | 0 | 0 | 0 | 0 | 0 |
| C4668T | 23 | 3 | 0 | 0 | 0 | 0 | 16 | 1 | 0 | 0 |
| C6539T | 8 | 6 | 0 | 0 | 1 | 0 | 1 | 0 | 0 | 0 |
| C7029T | 4 | 3 | 0 | 0 | 0 | 0 | 0 | 0 | 0 | 0 |
| C7819T | 10 | 3 | 0 | 0 | 1 | 0 | 0 | 0 | 0 | 0 |
| A9280G | 8 | 3 | 0 | 0 | 0 | 0 | 0 | 0 | 0 | 0 |
| C9803T | 16 | 3 | 0 | 0 | 1 | 0 | 1 | 1 | 0 | 0 |
| C9979T | 15 | 4 | 0 | 0 | 0 | 0 | 7 | 0 | 0 | 0 |
| G10465A | 11 | 5 | 0 | 0 | 0 | 0 | 0 | 0 | 0 | 0 |
| C13536T | 190 | 4 | 1 | 0 | 0 | 0 | 118 | 2 | 2 | 2 |
| G15906A | 11 | 4 | 0 | 0 | 0 | 0 | 0 | 0 | 0 | 0 |
| C18744T | 29 | 5 | 0 | 0 | 0 | 0 | 3 | 0 | 0 | 0 |
| C18877T | 595 | 3 | 1 | 2 | 0 | 0 | 30 | 9 | 7 | 0 |
| C19263T | 15 | 3 | 0 | 0 | 0 | 0 | 1 | 0 | 0 | 0 |
| G20398A | 7 | 3 | 0 | 0 | 0 | 0 | 0 | 0 | 0 | 0 |
| G20419T | 12 | 4 | 3 | 1 | 0 | 0 | 0 | 0 | 1 | 0 |
| G20887A | 10 | 4 | 0 | 0 | 2 | 0 | 0 | 0 | 0 | 1 |
| C21742T | 21 | 5 | 0 | 0 | 0 | 0 | 2 | 0 | 0 | 0 |
| C21855T | 18 | 9 | 0 | 0 | 0 | 0 | 1 | 0 | 0 | 0 |
| T21976C | 8 | 4 | 0 | 0 | 0 | 0 | 0 | 0 | 0 | 0 |
| A23975G | 4 | 4 | 0 | 0 | 0 | 0 | 0 | 0 | 0 | 0 |
| G25785T | 15 | 3 | 1 | 0 | 0 | 0 | 3 | 0 | 0 | 0 |
| C28629T | 20 | 4 | 0 | 0 | 0 | 0 | 1 | 4 | 1 | 2 |
| G28899T | 8 | 4 | 0 | 0 | 0 | 0 | 2 | 0 | 0 | 0 |
| C29719T | 6 | 3 | 0 | 0 | 0 | 0 | 2 | 1 | 0 | 0 |
| G29734C | 171 | 3 | 1 | 0 | 0 | 0 | 64 | 9 | 9 | 1 |
| G29779T | 13 | 4 | 0 | 0 | 0 | 0 | 5 | 0 | 0 | 0 |
| C23575T | 29 | 3 | 1 | 1 | 1 | 1 | 9 | 0 | 0 | 0 |
| C23587T | 17 | 3 | 0 | 1 | 0 | 0 | 6 | 0 | 0 | 0 |
| ATGA1605A | 484 | 9 | 1 | 0 | 2 | 0 | 306 | 108 | 0 | 1 |
| C29095T | 71 | 7 | 0 | 0 | 0 | 0 | 13 | 0 | 0 | 1 |
| C7164T | 12 | 2 | 0 | 0 | 0 | 0 | 5 | 1 | 0 | 0 |
| C619T | 21 | 2 | 0 | 0 | 0 | 0 | 0 | 0 | 0 | 0 |
| C25499T | 2 | 2 | 0 | 0 | 0 | 0 | 0 | 0 | 0 | 0 |

TABLE S2. **Summary of mutations for selected European countries as per May 26, 2020.** We include all mutations with respect to the Wuhan reference that appears at least three times in the Danish data. The last three mutations are included because they are relevant for the discussion in the section on Chains of mutations in Denmark.

### S3 Supplementary figures

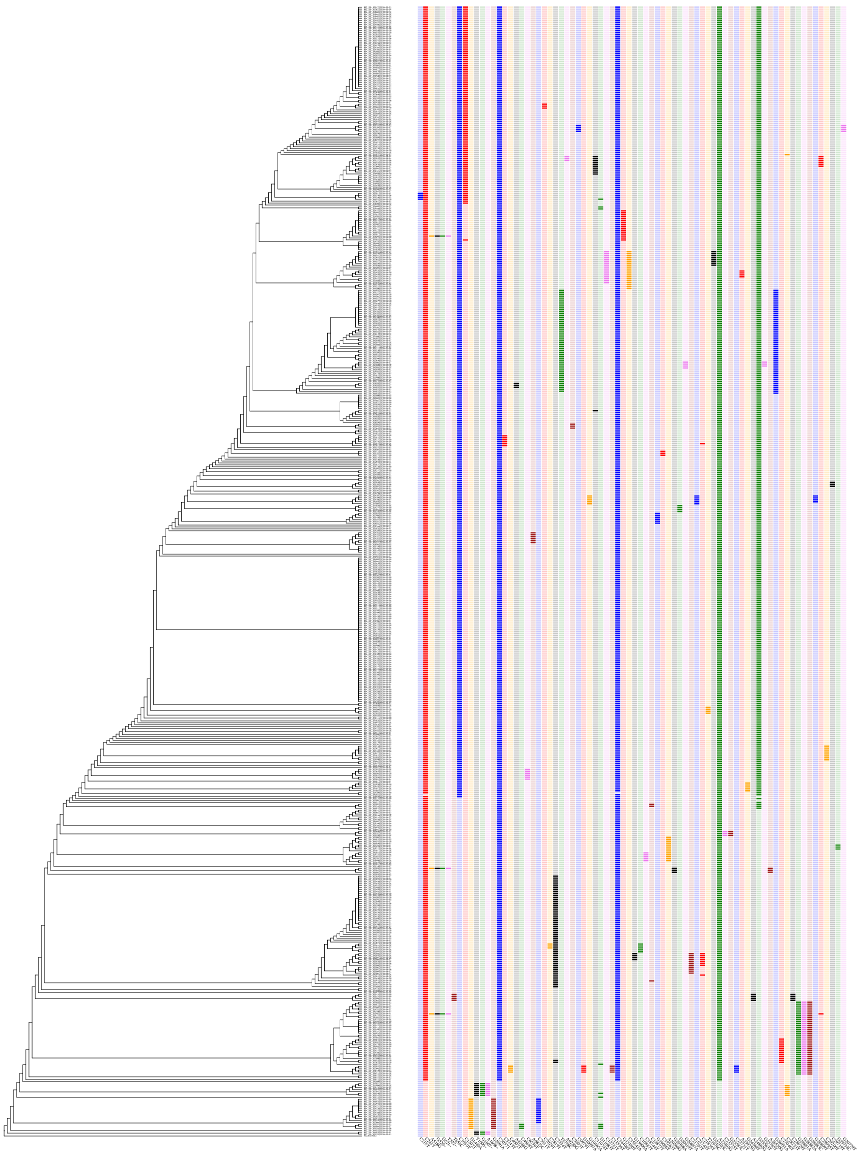

Fig S1. **Phylogenetic tree with Danish sequences.** Overview of the phylogenetic tree of Denmark on the left with the corresponding mutations on the right. Displayed are only mutations which happen at least thrice. The plot shows how about 70% of the sequences have the haplotype A2a2a (corresponding to mutations C241T, C3037T, A23403G, C14408T, G25563T, C1059T) and that 103 have the further mutation C1302T. Note that the mutation A518G with the co-occurring deletion 520-522 is equivalent to the deletion 518-520 which is listed in [31] as can easily be seen by inspecting the Wuhan sequence.

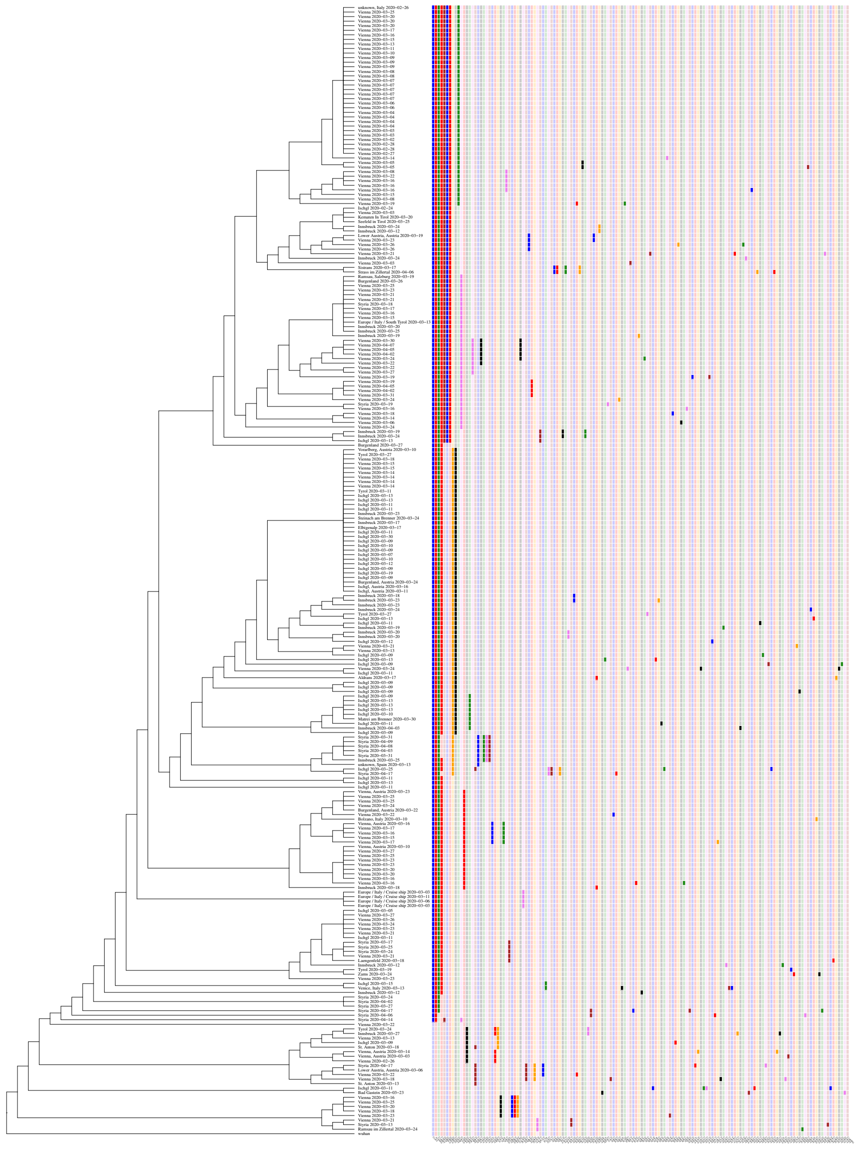

FIG S2. **Phylogenetic tree with Austrian sequences.** Austrian sequences with location metadata and mutations with respect to the Wuhan standard reference.

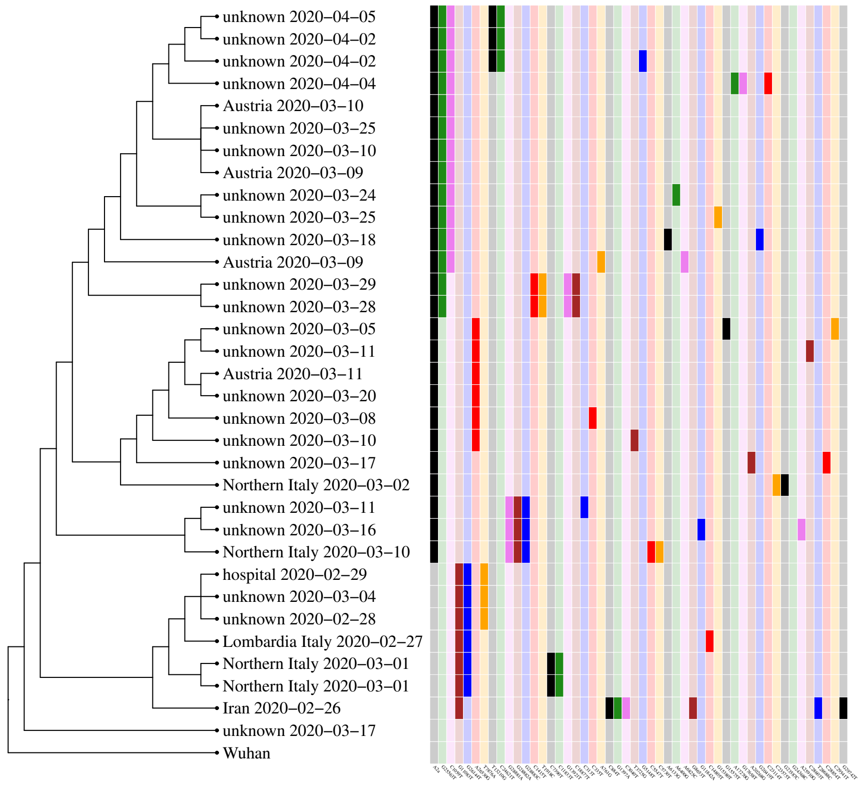


FIG S3. **Phylogenetic tree with Norwegian sequences.** The tree shows the Norwegian sequences with travel history metadata and mutations with respect to the Wuhan standard reference.
